# PhaMers identifies novel bacteriophage sequences from thermophilic hot springs

**DOI:** 10.1101/169672

**Authors:** Jonathan Deaton, Feiqiao Brian Yu, Stephen R. Quake

## Abstract

Metagenomic sequencing approaches have become popular for the purpose of dissecting environmental microbial diversity, leading to the characterization of novel microbial lineages. In addition of bacterial and fungal genomes, metagenomic analysis can also reveal genomes of viruses that infect microbial cells. Because of their small genome size and limited knowledge of phage diversity, discovering novel phage sequences from metagenomic data is often challenging. Here we describe PhaMers (<u>Phage</u> *k*-<u>Mers</u>). a phage identification tool that uses supervised learning to classify metagenomic contigs as phage or non-phage on the basis of tetranucleotide frequencies. a technique that does not depend on existing gene annotations. PhaMers compares the tetranucleotide frequencies of metagenomic contigs to phage and bacteria references from online databases. resulting in assignments of lower level phage taxonomy based on sequence similarity. Using PhaMers. we identified 103 novel phage sequences from hot spring samples of Yellowstone National Park based on data generated from a microfluidic-based mini-metagenomic approach. We analyzed assembled contigs over 5 kbp in length using PhaMers and compared the results with those generated by VirSorter, a publicly available phage identification and annotation package. We analyzed the performance of phage genome prediction and taxonomic classification using PhaMers. and presented putative hosts and taxa for some of the novel phage sequences. Finally. mini-metagenomic occurrence profiles of phage and prokaryotic genomes were used to verify putative hosts.

## Introduction

Bacteriophages (phages) are viruses that infect microorganisms such as bacteria or archaea. Phage play important roles in microbial communities including gene duplication [1] and lateral gene transfer [2], and are also the most abundant biological entities on the planet at an estimated 10^31^ viral particles [3]. Despite this abundance, our understanding of phage diversity is limited to thousands of partial or completed sequences. Before the advent of high-throughput sequencing, our understanding of phage genomics was limited to lineages that could be cultured in plaques [4]. Although metagenomics alleviates this limitation [5], the typically small size of phage genomes and the lack of a universal marker gene complicate the identification of phage genomic sequences [6].

Common approaches of phage identification use Hidden Markov Models (HMM) based gene annotation to search for coding regions that are homologous to known viral genes [7], including “hallmark genes” such as the terminase large and small subunits, major capsid, coat, tail, and portal proteins [8]. Approaches based on hallmark genes perform well for identifying genomes closely related to known phage sequences but may fail at identifying phage sequences with little known genetic homologues. A less common approach involves considering the frequencies of short oligonucleotides of length *k* (*k*-mers) [9]. Frequencies of *k*-mers vary among species, and can be used to predict phylogeny and taxonomy. For instance, *k*-mer frequencies of a phage’s genome are predictive of the phage’s host [10] [4] [11], and that *k*-mer frequencies in 16S rRNA sequences can be used to predict bacterial taxonomy [12] [13]. A k-mer frequency based approach is advantageous when no high-similarity alignments exist. Such an approach also reduces computational expense compared to alignment-based methods [14] [15].

Here we present the characterization and application of PhaMers (<u>Phage</u> *k*-<u>Mers</u>), a novel phage identification algorithm that uses tetranucleotide frequency and supervised learning algorithms to identify viral sequences from metagenomic sequencing data. We compared the results of PhaMers to those from VirSorter, an automated phage identification pipeline based on alignment with hallmark genes [8]. We applied both PhaMers and VirSorter to metagenomic contigs from three hot spring locations in Yellowstone National Park (YNP), resulting in 913 novel bacteriophages, out of which 103 sequences were also supported by annotated genes. We used Joint Genome Institute’s Integrated Microbial Genomes & Microbiomes (IMG/ER) pipeline [16] to generate functional annotations for all phage sequences. In addition, we explored the relationship between tetranucleotide frequency and phage taxonomy both in a reference dataset and among novel phages from YNP.

## Results and discussion

### Tetranucleotide frequency differentiates phage taxonomy

Tetranucleotide frequency can be used to group prokaryotic genomes and to assign taxonomy [17]. Due to viral host specificity, we argue that tetranucleotide frequencies can also serve as a statistic to distinguish phage genomes [11] [18] [19]. To assess relationships in tetranucleotide frequencies among known phage sequences, we collected 2255 phage genomes from RefSeq in October 2015 belonging to various taxonomies and having different genome lengths (S1 Fig and S1 Table), calculated tetranucleotide frequencies, and visualized phage genome relations in two dimensions using t-Distributed Stochastic Neighbor Embedding (t-SNE) [20] (Fig 1a). We then clustered phage genomes using DBSCAN [21], and observed enriched taxa within clusters (Fig 1b). 60% of the phage genomes were assigned to clusters (S2 Table). We noticed that varying clustering parameters reveal similarities between viral taxa. Strict cluster assignments resulted in the increase of cluster purity under Baltimore classification. More liberal assignments, on the other hand, resulted in clusters with multiple closely related phage phylogenies (S2 Fig). For instance, *k*-means clustering with a less stringent *k* (*k*=40) resulted in the assignment of dsDNA *Cellulophaga phage phiSM*, dsDNA *Lactococcus phage 936 sensu lato*, and *Skunalikevirus* to the same cluster (cluster 10 in S3 Table), indicating that phage of these taxa have similar genomic tetranucleotide frequencies, and may be closely related.

**Fig 1.**
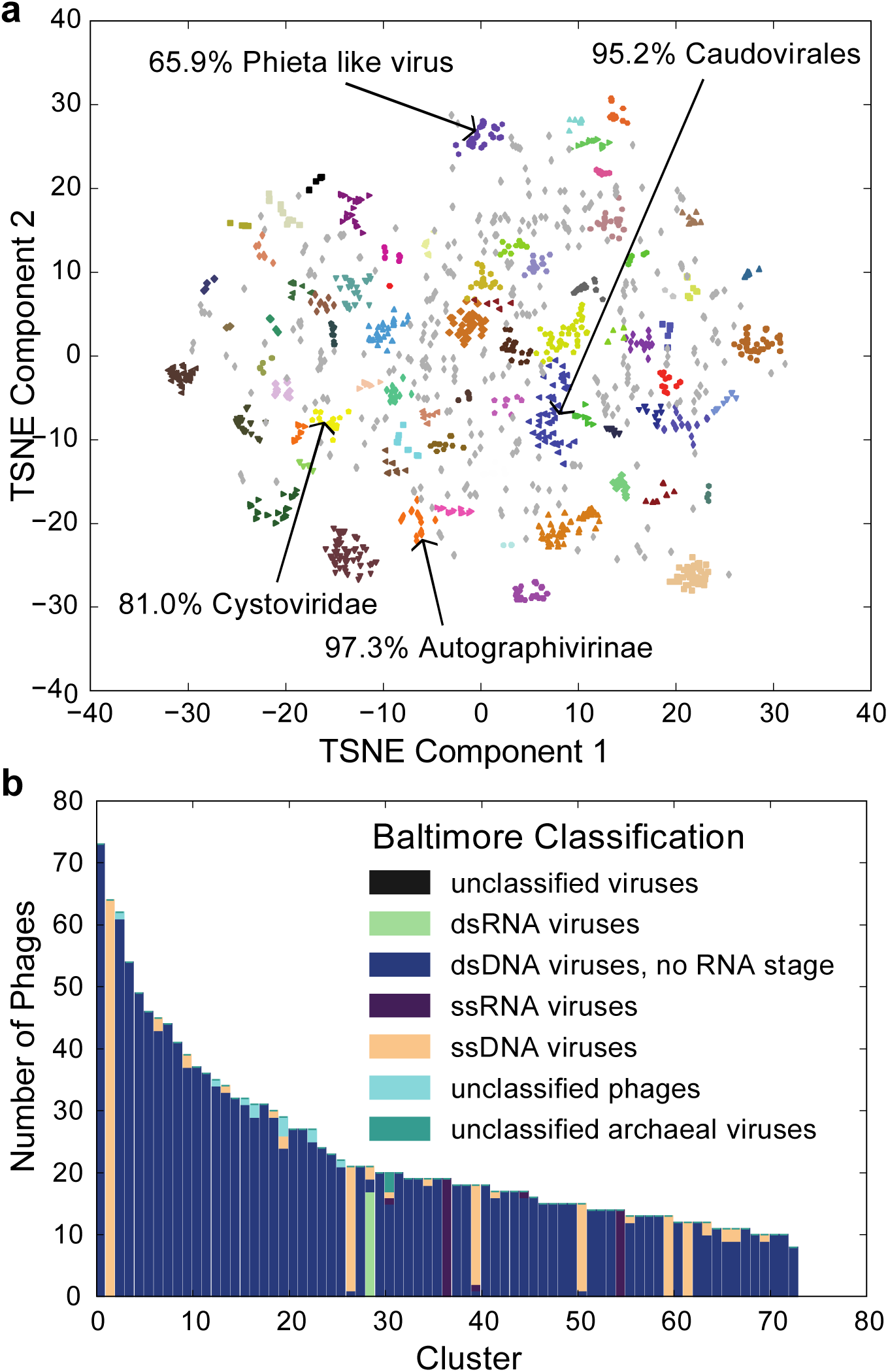
RefSeq phage tetranucleotide characteristics and taxonomy. **a**, t-SNE representation based on tetranucleotide frequency of 2255 phage sequences from RefSeq. Clusters assigned with reduced dimensionality (2D) embedded tetramer frequency vectors and clustered using DBSCAN (epsilon = 1.5, min points per cluster = 10). Some clusters enriched with phage of a single taxon are labeled with percentages denoting the proportion of phage sequences in that cluster belonging to that enriched taxon. A cluster is considered enriched with a taxon if the proportion of phage sequences belonging to that taxon in the cluster is greater than 50% and the enriched abundance is statistically significant compared to background abundance in the reference dataset, as tested using Pearson’s chi-squared test. **b**, Compositions of taxa for phage sequences assigned to the clusters shown in Fig 1a at the Baltimore Classification depth.

Because most single stranded RNA (ssRNA) viruses belong to the genera *Allolevirus* and *Levivirus*, generally classified as *Enterobacteriophge MS2* and *Enterobacteriophage Qß* [22], they are primarily classified into two pure clusters (Fig 1). Using *k*-means clustering with *k*=40 also resulted in the assignment of both taxa to cluster 8 (S2b,c Fig and S3 Table), indicating *Allolevirus* and *Levivirus* each have signature tetranucleotide frequencies more similar to each other than to most other phages. Single stranded DNA phage is another taxon which tends to forms enriched clusters. The biggest cluster is enriched for the well-characterized 5.4 kbp *Enterobacteria phage phiX174 sensu lato*. The second largest of these clusters is enriched for the 5.5 kbp *Enterobacteria phage G4 sensu lato*. Tetranucleotide frequency based clustering also enriches for double stranded DNA (dsDNA) phage families such as *Siphoviridae*, *Myoviridae*, and *Podoviridae*, which are under the broader family of *Cystoviridae* (Fig 1). Previous work has shown that alignment free methods are capable of producing accurate viral taxonomic classifications [23] [24]. Clusters of phage genomes enriched for a single phage taxon, therefore, indicate that tetranucleotide frequency based genome classification is an effective phage differentiation scheme.

### PhaMers uses machine learning techniques to identify phage sequences from metagenomic contigs

In addition to differentiating phage sequences from each other, tetranucleotide frequencies can differentiate phage from prokaryote sequences. This feature is useful since the abundance of prokaryotic contigs from metagenomic datasets often hampers discovery of novel viral genomes. Hence, we sought to understand how tetranucleotide frequencies could best distinguish phage sequences from prokaryotic sequences and developed a machine-learning algorithm that compares tetranucleotide frequencies of unknown sequences to those of known phage, bacterial and archaeal sequences (S3 Fig). Reference phage genomes from RefSeq (S1 Table) and bacterial and archaeal genomes from GenBank (S4 Table) were used. Testing a set of machinelearning algorithms and their combinations using 20-fold cross validation on reference phage genomes,we found that a linear combination of KNN and a cluster centroid proximity metric (S1 Text and S4 Fig) performed best when considering both area under the curve (AUC = 0.992) and true positive rate (TP = 91%) (Fig 2a). We decided to use this combined approach for phage identification in PhaMers. PhaMers scores predicted phage sequences with positive values and non-phage sequences with negative values (Fig 2b). Well-studied and heavily represented phage sequences in the dataset were more likely to be classified correctly, yielding 91.8% sensitivity, *99.3%* specificity, and 99.2% positive predictive value (PPV).

**Fig 2.**
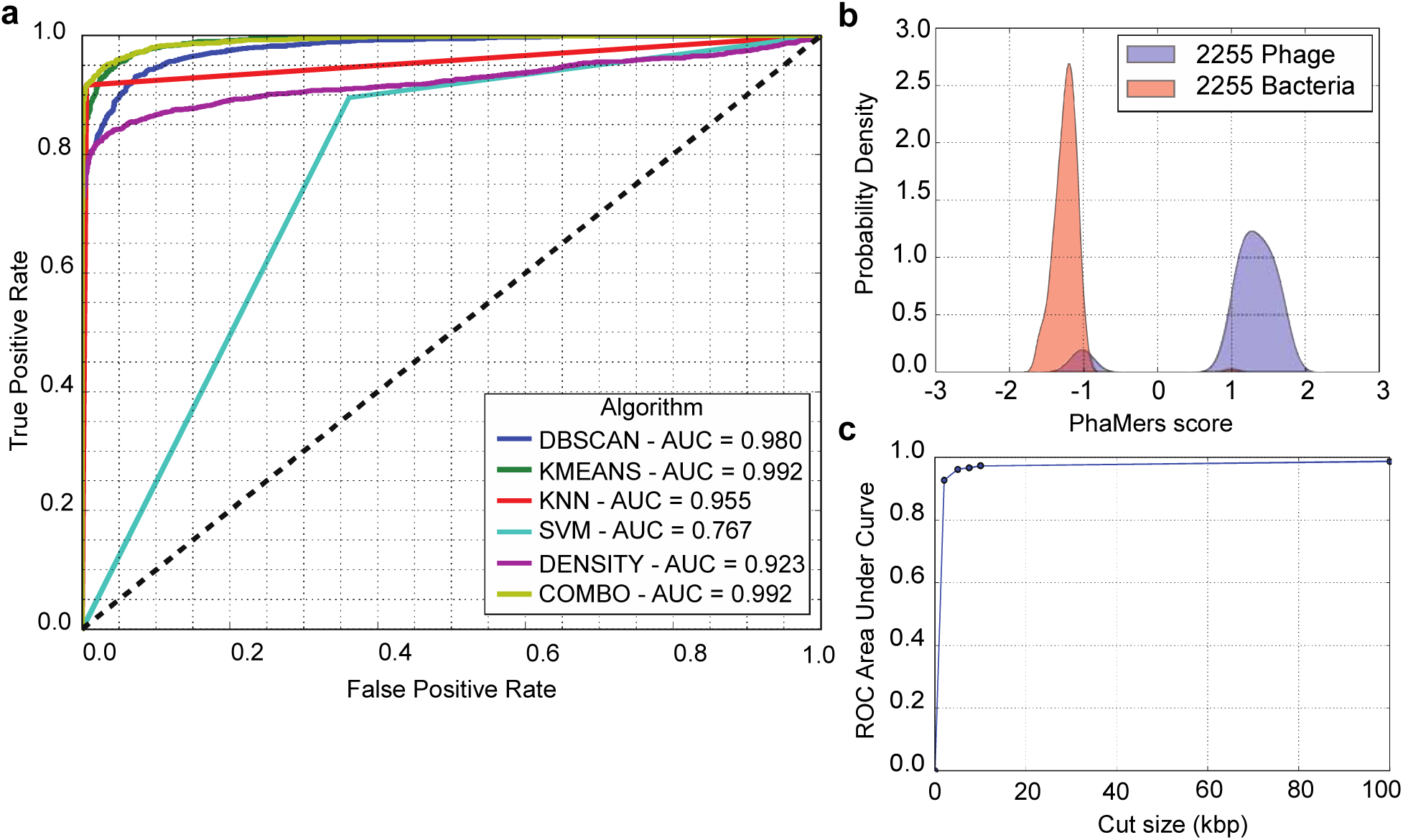
*N*-fold cross validation of PhaMers scoring algorithm. **a**, Receiver operator characteristic (ROC) curves for various supervised learning algorithms used on the reference datasets of phage and bacterial sequences in 20-fold cross validation. Feature vectors are tetranucleotide frequencies calculated from the 2255 phage and 2255 bacterial sequences in the reference dataset. “DBSCAN” and “KMEANS” scoring algorithms apply the method described in S1 Text to calculate a metric of proximity to the nearest clusters of phages and bacteria assigned by DBSCAN and *k*-means clustering, respectively. “DENSITY” stands for a custom algorithm that uses Kernel Density Estimation to approximate the probability density of phage and bacteria data points, and then gives a score as the log-ratio of the two probabilities. “KNN” represents a K-Nearest-Neighbors algorithm, and “SVM” represents a Support Vector Machine approach. The “COMBO” algorithm is a linear combination of K-Nearest Neighbors and “KMEANS”. This scoring algorithm, described in S1 Text, was chosen for use in subsequent applications of PhaMers. The “COMBO” algorithm was chosen over “KMEANS” because although both had AUC = 0.992, the true positive rate of “COMBO” was 91% as compared to 84% for “KMEANS”, when the classification threshold was set to zero. **b**, Distributions of PhaMers scores, given by “COMBO”, for 2255 phage and 2255 bacterial genomes, as calculated with 20-fold cross validation. The small blue population centered at score = ࢤ1 are the phages in the datasets which were misclassified as bacteria (false negatives). There is also a smaller population of misclassified bacteria shown at score = 1. **c**, Predictive performance (AUC) of PhaMers as a function of reference genome length. The reference datasets of phage and bacterial genomes were cut randomly to sizes of 2.5 kbp, 5 kbp, 7.5 kbp, 10 kbp, and 100 kbp. Predictive performance, shown on the y-axis, is given by the area beneath the ROC, which drops as sequences were cut to lengths shorter than 5 kbp. Dataset of bacteria used for this analysis are included in S6 Table.

When classifying reference sequences that had been omitted from the training set, PhaMers correctly classified phage sequences belonging to *Enterobacteria phage T4*, T4-like and T7-like viruses, *Propionibacterium phage*, and *Lactococcus phage ASCC*. Misclassified phage sequences (negative scores) were generally from *unclassified Siphoviridae*, *unclassified Podoviridae*, and *unclassified Myoviridae*. Misclassification occurred because there were few similar relatives in the dataset. To discount the effect of heavily represented phages on performance estimation, we performed cross validation with a reduced dataset of phages containing only one member from each taxonomy. Tetramer frequencies distinguished this set of phage sequences from bacteria with only marginally decreased accuracy (S5 Fig). We also assessed how sequence length affected PhaMers’ performance by performing 20-fold cross validation with random subset from the test dataset. We observed that the area under the curve (AUC) of the receiver operator characteristic (ROC) dropped most significantly as sequence length decreased below 5 kbp (Fig 2c), demonstrating that 5 kbp is a useful contig length cutoff.

### PhaMers identifies putative phage sequences from Yellowstone hot springs

We analyzed two sets of assembled metagenomic contigs longer than 5 kbp with VirSorter and PhaMers and visualized the results (Fig 3a). These datasets were prepared using a microfluidic-based mini-metagenomic method on two hot spring samples from Bijah and Mound Springs of the Yellowstone National Park (Materials and methods) [25]. PhaMers identified 836 of the 5594 contigs longer than 5 kbp as potential phage with positive scores. VirSorter identified 30 putative phage contigs from the Bijah Spring sample and 108 from the Mound Spring sample. Six phage contigs from the Mound Spring dataset were identified as potentially prophage (Fig 3b). Of the 89 contigs in the Mound Spring dataset identified as potentially phage-like by VirSorter, 52 were given positive PhaMers scores. Of the 23 contigs in the Bijah Spring dataset identified as potentially phage-like by VirSroter, 10 were given positive scores by PhaMers. In the Bijah Spring and Mound Spring datasets, 261 and 575 contigs, respectively, received positive PhaMers scores but were not labeled as phage-like by VirSorter. In total, 1663 and 3036 contigs, respectively, were not labeled as phage-like by either VirSorter or PhaMers. Although some predictions made by PhaMers and VirSorter agree, many contigs received positive PhaMers scores but were not labeled as phage-like by VirSorter, indicating that PhaMers may identify phage genomes that VirSorter misses. While the nearly 700 such contigs may represent previously unidentified and highly divergent phages, verifying their biological role will require substantial follow up and thus for the remainder of this study we focus on those sequences which are concordant between the algorithms.

**Fig 3.**
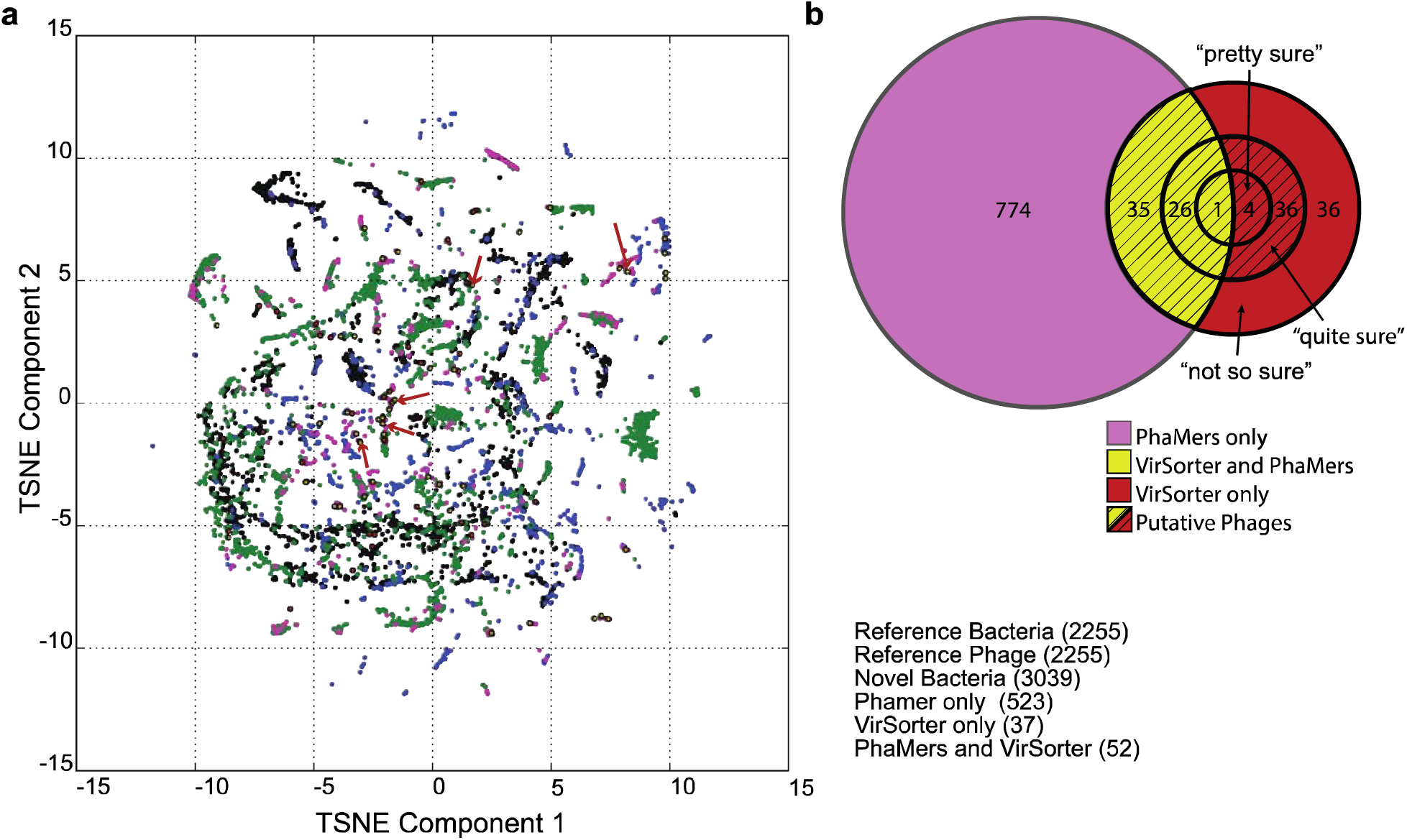
PhaMers and VirSorter analysis overview. **a**, t-Distributed Stochastic Neighbor Embedding (t-SNE) representation of tetramer frequency vectors of phage (blue) and bacterial (black) references in conjunction with contigs longer than 5 kbp from the Mound Spring dataset. Points representing contigs are colored according to whether they are designated as viral by VirSorter (Red), PhaMers (purple), both (yellow), or neither (green). Red arrows indicate contigs identified as phages which were assigned a taxonomic classification based on their proximity to clusters of phages enriched for a single taxon. **b**, Venn diagram showing the number of contigs from Mound Spring and Bijah Spring which were classified as viral by PhaMers and VirSorter. Contigs considered to be putative phages (shaded) were those which were either classified as viral with high confidence (“quite/pretty sure”) by VirSorter, or were classified with low confidence (“not so sure”) but which received a positive PhaMers score.

To deepen our analysis of these novel phage sequences, we used t-SNE to visualize tetranucleotide frequencies of all contigs combined with phage and bacterial reference sequences (Fig 3a and S6 Fig). Many contigs form tight clusters, likely originating from closely related bacterial genomes. Although PhaMers identified greater viral diversity than VirSorter, many contigs only contained unannotated open reading frames, precluding any biological interpretation. In order to examine annotated gene functions, we focused on a set of putative phage contigs predicted by VirSorter and PhaMers that lie in proximity to clusters of known phage genomes (19 contigs from Bijah Spring and 83 contigs from Mound Spring). We also analyzed a single contig of 35 kbp from a third metagenomic dataset generated from a Mammoth Geyser Basin sample. This contig was identified as a phage sequence by PhaMers and BLASTP, but not VirSorter, demonstrating PhaMers’s ability to uncover phage sequences that other methods miss. Contig annotations using the Integrated Microbial Genomes and Microbiomes (IMG) gene annotation pipeline (Materials and methods) further supports PhaMers and VirSorter predictions (S5 Table).

To predict phage taxonomy from tetranucleotide frequencies, we used *k*-means to cluster known phage tetramer frequency vectors with those of novel phages. We looked for novel phage contigs assigned to clusters enriched with reference phage sequences of a single taxon. Clusters were labeled as enriched for a taxon if members of that taxon constituted more than half of the cluster and their prevalence was significantly greater than the proportion that the taxon was represented in the reference dataset (S1a Fig). A contig meeting these criteria and lying within a standard deviation of the mean cluster silhouette value were assigned the taxon of that cluster. Of the 24 contigs (five from Bijah Spring, 18 from Mound Spring, and one from Mammoth Geyser Basin) that met these criteria, most were assigned to clusters enriched for *Siphoviridae*, *Podoviridae*, and *Myoviridae* (S5 Table Column I and Fig 3a red arrows). These three families of dsDNA phages belong to the order *Caudovirales* and are differentiated by their tail morphology [26]. *Podoviridae* have short non-contractile tails while *Myoviridae* and *Siphoviridae* have long tails, contractile for the former and non-contractile for the ladder [26].

We discovered several clear phage sequences due to predicted taxonomy and gene phylogeny. A 13,504 base pair phage contig (Contig 1753) from Mound Spring (category 2 “quite sure” prediction by VirSorter and 1.14 by PhaMers) is similar in tetranucleotide frequency to *Siphoviridae* and contains phage tail and portal proteins typically associated with *Siphoviridae* (Fig 4a-c). However, 15 of the 19 putative proteins are homologous to genes from the bacterial lineage *Clostridiales*. In addition, analyzing occurrence patterns, we notice that contig 1753 is only present in one mini-metagenomic subsample. We observe similar occurrence patterns across another set of 9 contigs also assigned to *Clostridiales* (Fig 4d). None of the other genomes appear only in the same mini-metagenomic subsample, suggesting that contig 1753 is more likely a *Siphoviridae* that infects *Clostridiales*, although we cannot tell if the *Siphoviridae* fragment is inserted into a *Clostridiales* genome because of its rare occurrence. Another phage contig (Contig 677) 20,664 base pairs in length (category 2 by VirSorter and 0.97 by PhaMers) is enriched (33/36) for genes homologous to the thermophilic bacterium *Hydrogenobacter*, in the phylum *Aquificae* (Fig 4d, e). The enrichment for predicted phage genes homologous to *Hydrogenobacter* is an indication that *Hydrogenobacter* is a natural host. This hypothesis is validated by the similar and abundant occurrence patterns between the phage contig and a partial *Hydrogenobacter* genome from the same environment (*p* < 10^−12^, Fisher’s exact test) (Fig 4d). While it may be possible that an infection is so prevalent that the phage is observed to co-occur with its host in a large fraction of the hosts, it seems more likely that high co-occurrence statistics are a sign that the phage is either incorporated into the host genome as a prophage or exists in the host as a plasmid. For example in this case the co-occurrence pattern suggests that the contig is likely integrated in the genome of its host or exists as a plasmid because 83% of the sub-samples containing the *Hydrogenobacter* genome also contain parts of the viral genome.

**Fig 4.**
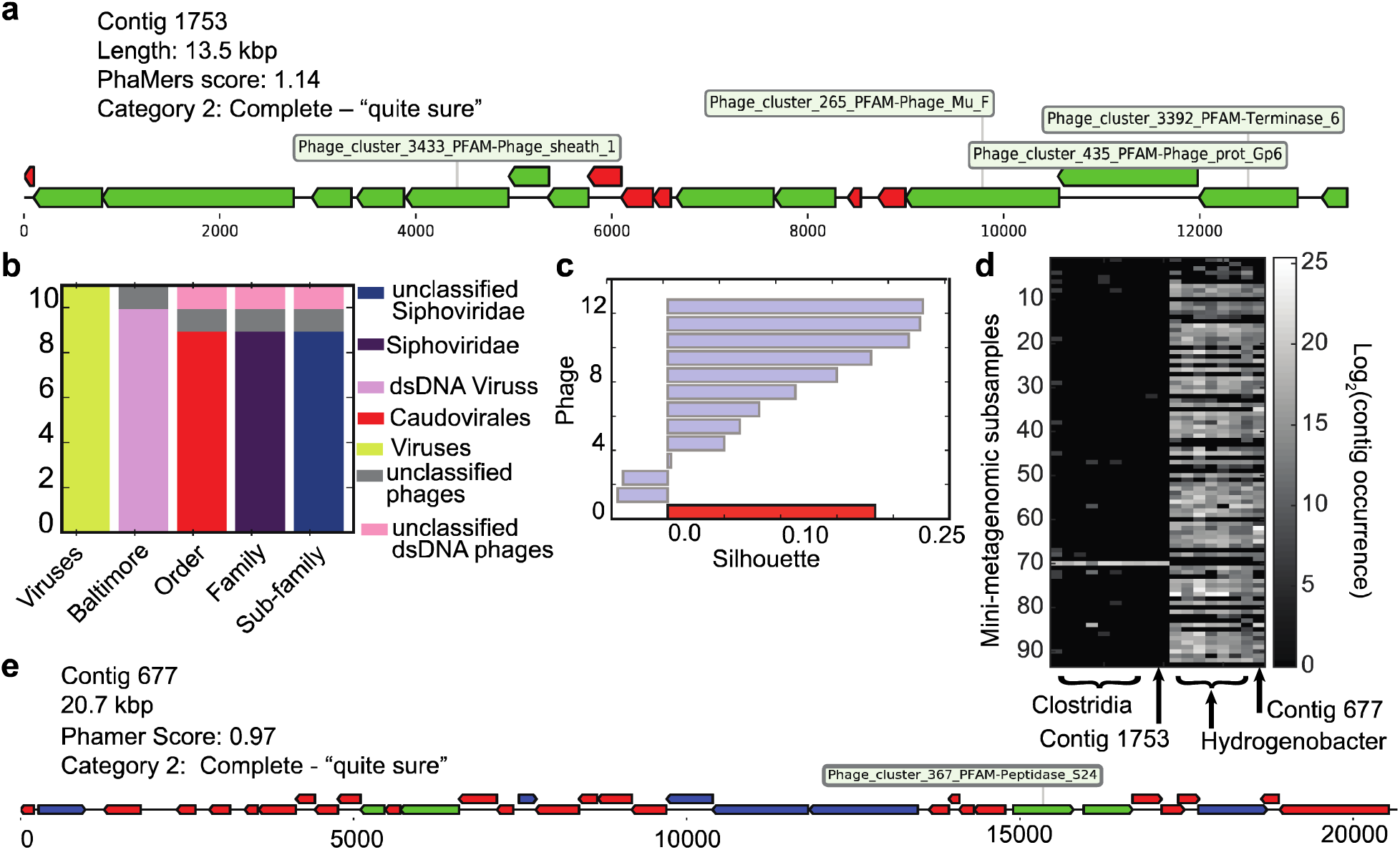
Selected novel phage contigs. **a**, Diagram of 13.5 kbp contig (Contig 1753) from Mound Spring showing putative coding regions identified by VirSorter. Green sections represent genes with viral function, blue sections represent genes with known but non-viral annotations, and red sections represent hypothetical proteins. **b**, Bar graph representing the composition of taxa for phages in the cluster to which contig 1753 was assigned by *k*-means (*k*=86) on the basis of tetranucleoitde frequencies. **c**, Cluster silhouette values for reference phage sequence (blue) and contig 1752 (red) assigned by *k*-means clustering. **d**, Contig occurrence patterns across 93 mini-metagenomic subsamples. Contig 1753 is plotted with related Clostridia contigs with similar occurrence patterns. Contig 677 is plotted along with a close by cluster of *Hydrogenobacter* contigs. Brightness represents base 2 logarithm of the contig occurrence within each subsample, defined as the number of basepairs covered by at least 1 read in that subsample. **e**, Diagram of contig 677 showing VirSorter annotations. Green sections represent genes with viral function, blue sections represent genes with known but non-viral annotations, and red sections represent hypothetical proteins.

Finally, in the Mammoth Geyser Basin sample, we characterized a single novel viral contig of length 35,211 base pairs (S6 Fig). This contig was not identified as a phage by VirSorter but was given a score of 0.90 by PhaMers. NCBI BLAST revealed similarities at both nucleotide and protein level to the thermophilic archaeal phage genera *Sulfobus filamentous* and *Acidianus filamentous*, belonging to the *Betalipothrixvirus* genus of the family *Lipothrixviridae* [27]. Interestingly, this contig was assigned on the basis of tetranucleotide frequencies to a cluster enriched with *Lactococcus phage 936 sensu lato*, which were *unclassified Siphoviridae*. Protein blast revealed a 596 amino acid putative protein with 73% identity to a Holiday junction branch migration helicase from *Acidianus filamentous virus* 9, as well as a 563 amino acid putative protein with 72% identity to a helicase from *Acidianus filamentous virus* 9. Another 1038 amino acid putative protein had 48% identity to a conserved hypothetical protein of *Acidianus filamentous virus 3* [28], not appearing in the *Sulfobus filamentous* genome [27]. These features indicate that as a new phage in the *Betalipothrixvirus* genus, this contig has shorter phylogenetic distance to *Acidianus filamentous* than to *Sulfobus filamentous*. Taken together, PhaMers offers a new method to discover phage contigs from metagenomic datasets that may be missed by gene-based methods.

## Conclusion

The large numbers of recent metagenomic studies have generated exponentially increasing environmental sequencing data representing large prokaryotic and viral diversity. Although significant progress has been made in mining prokaryotic genomes from metagenomic datasets, finding phage genomes is still difficult, partly due to the lack of universal marker genes among phage genomes. The tetranucleotide frequency and machine learning aspects of the PhaMers algorithm enable identification of phage contigs without depending on protein homology, which is valuable because most viral genes are poorly annotated. In this study, PhaMers identified genomes missed by VirSorter, demonstrating that PhaMers complements existing approaches for phage discovery. In addition to phage contig identification, PhaMers permits phage classification and the identification of potential host by assessing proximity of nearby phage or prokaryotic genome clusters in the tetranucleotide frequency space. Combined with mini-metagenomic co-occurrence patterns, our computational method has the potential to differentiate between a phage infection and bacterial cells that carry phage genomes. PhaMers requires less computing resources compared to methods that perform sequence alignments because distances are computed only in the tetranucleotide frequency space. Although PhaMers’s performance is affected by the comprehensiveness of reference datasets, most of which are dominated by few, well-characterized phage taxonomies, we have taken steps to mitigate such effects by using a reduced reference subset during cross validation. As the number of metagenomic datasets continues to increase, more phage genomes will be discovered, leading to more comprehensive reference phage databases. As phage databases grow to include a more comprehensive representation of global diversity, PhaMers analysis could represent a scalable, computationally light method for rapidly assigning phage taxonomic classifications.

## Materials and methods

### Sample collection

Environmental samples used in this study were collected from two separate hot springs from Yellowstone National Park under permit number YELL-2009-SCI-5788: sediments of the Bijah Spring in the Mammoth Norris Corridor and Mound Spring in the Lower Geyser Basin region. Samples were placed in 2 mL tubes and soaked in 50% ethanol onsite. Samples were spaced in 2 mL tubes without any filtering and soaked in 50% ethanol onsite. Upon returning, samples were transferred to ࢤ80°C for long term storage. Biosample information can be found from Joint Genomes Institute’s GOLD system under Gb0114344 and Gb0114821 respectively.

### Sample preparation and sequencing

Environmental samples were processed using the microfluidic-based mini-metagenomic protocol performed on a commercially available Fluidigm C1 Auto Prep IFC (Integrated Fluidic Circuit)[25], with scripts and detailed protocols available on Fluidigm’s ScriptHub (https://www.fluidigm.com/c1openapp/scripthub/script/2017-02/mini-metagenomics-qiagen-repli-1487066131753-8). Steps performed on the automated microfluidic platform included cell partition, cell lysis, and genomic DNA amplification using MDA (Multiple Displacement Amplification). During this process, both intracellular and environmental phage DNA were amplified. Amplified genomic DNA was harvested into a 96-well plate. The concentration was quantified independently using the high sensitivity large fragment analysis kit (AATI) and adjusted to 0.1–0.3 ng/μL, the input range of the Nextera XT library prep pipeline. Nextera XT V2 libraries (Illumina) were made and sequenced on the Illumina NextSeq (Illumina) platform using a 2x150 bp runs. Sequencing reads were filtered and assembled according to the methods described by Yu et al. [25]. Assembly was performed via SPAdes V3.5.0 with kmer values of 33, 55, 77, and 99. Contigs longer than 5 kbp were retained for phage analyses.

### PhaMers classification of metagenomic contigs

PhaMers’ scoring algorithm was used to score assembled metagenomic contigs greater than 5 kbp. PhaMers uses BioPython to parse fasta formatted files of assembled contigs, and tabulates tetranucleotide frequencies before scoring. Results are written to file and are used for subsequent analysis. To assign putative taxonomic classifications to phage, we used *k*-means (*k*=86) to cluster the tetranucleotide frequencies of each putative phage with those of the phages from the reference data set, and examined the taxonomic composition of the phage sequences in the cluster that the contig was assigned to. We considered a contig assigned to a taxonomically enriched cluster if phage from a single taxon compose its cluster at a proportion greater than 50% and statistically significantly greater than the proportion that that taxa represents in the entire dataset. We considered it evidence of a putative phage’s taxonomic classification if a phage was assigned to an enriched cluster and if its cluster silhouette score was within one standard deviation of the mean cluster silhouette scores of the reference phage in the cluster.

### Reference database generation

The reference dataset of genomic phage sequences was assembled using the Phage available on RefSeq in October of 2015. A complete list of all viral accession numbers made available on NCBI was downloaded and used to find accession numbers for all viruses that infect bacteria or archaea (S1 Table). This set of accession numbers was used to access and compile a set of all phage sequences in fasta format, subsequently used for tetranucleotide frequency analysis. The reference dataset of bacterial genomic sequences was generated from genomic assemblies available on GenBank (S4 Table). Bacterial species were selected at random from the subdirectories at ftp.ncbi.nlm.nih.gov/genomes/genbank/bacteria/, and the latest genomic assembly fasta files were used for analysis in PhaMers. Tetranucleotide frequencies were calculated for all genomic fragments within each contig file, and a list of bacterial accession number a tetranucleotide counts.

### PhaMers classification of known phage and bacterial genomes

To verify PhaMers’ predictive discrimination between phage and non-phage genomic sequences, we selected a *k*-mer length of 4 (tetranucleotide) and counted occurrences of each of the 256 tetramers in the 2255 phage and 2255 bacterial genomes of our reference dataset. We then normalized each tetranucleotide count vector by the total number of tetramers counted to produce tetranucleotide frequency vectors, thereby discrediting differences in vectors due to sequence length. We then performed 20-fold cross validation on different scoring algorithms, wherein we divided both the phage and bacterial datasets in to subdivisions and scored each subdivision by using the remaining nineteen subdivisions as training data.

We tested the following supervised learning algorithms: Support Vector Machine, Kernel Density Estimation, K-Nearest Neighbors (KNN), and Nearest Centroid, and linear combinations of results from each (Fig 2a). KNN was chosen as the primary classifier because it performed with lowest false positive rate while maintaining >90% sensitivity. We varied the parameter specifying number of neighbors used in KNN classification (K) from 3 to 20 during cross validation on our reference datasets. Increasing K yielded marginally decreased performance, hence informing our choice of K=3. To add additional information into the final PhaMers score, we took the initial classification by KNN to be a ࢤ1 (non-phage) or 1 (phage) and added to it a parameter between ࢤ1 and 1 that quantified the proximity of a point to phage clusters, and distance away from bacterial clusters. (S1 Text and S4 Fig) This was chosen because this algorithm performed well on its own, and quantifies relative distances to large groups of reference data, whereas KNN classified based on more local data points.

To study the relationship between tetranucleotide frequencies and phage taxonomy, we reduced the dimensionality of this reference phage tetramer frequency vectors from 256 to two using t-SNE (Fig 1a). We then clustered the reduced dimensionality tetramer frequency vectors using density-based spatial clustering DBSCAN, [21] and quantified the prevalence of each taxa in each cluster.

### VirSorter analysis of metagenomic contigs

Samples were analyzed using the VirSorter 1.0.3 phage identification pipeline available through the iPlant Discovery Environment on the iPlant collaborative website, made available by CyVerse. (https://de.iplantcollaborative.org/de/) VirSorter used all bacterial and archaeal virus genomes in Refseq, as of January 2014 for the analysis of both metagenomic samples.

### Annotation of putative phage contigs

Contigs were uploaded to JGI’s Integrated Microbial Genomes Expert Review online database (IMG/ER). Annotation was performed via IMG/ER [31]. Briefly, structural annotations were performed to identify CRISPRs (pilercr), tRNA (tRNAscan), and rRNA (hmmsearch). Protein coding genes were identified with a four ab initio gene prediction tools: GeneMark, Prodigal, MetaGeneAnnotator, and FragGeneScan. Functional annotation was achieved by associating protein coding genes with COGs, Pfams, KO terms, and EC numbers. Phylogenetic lineage was assigned to each contig based on gene assignment. Annotations can be found under genome IDs 3300006068 and 3300006065.

### PhaMers development

Code and algorithms used by PhaMers were tested in MATLAB and implemented in the Python 2.7 programming language. Python 2.7 was used to write scripts for parsing of VirSorter and IMG output files and for integration with PhaMers data. All PhaMers scripts are available at https://github.com/jondeaton/PhaMers. The Python library Matplotlib was used for plot generation. We automated the visualization of contig gene annotations, with a Python package named dna_features_viewer 0.1.0 (pypi.python.org/pypi/dna_features_viewer/0.1.0)

## Acknowledgments

The authors of this work would like to acknowledge members of the Quake Lab Sequencing Facility: Norma Neff, Jennifer Okamoto, Gary Mantalas, and Ben Passarelli. The authors would also like to acknowledge generous support from the DOE JGI’s Emerging Technologies Opportunities Program (ETOP), John Templeton Foundation, Stanford Graduate Fellowship, NSF GRFP, and the Stanford Bioengineering REU program. The work conducted by the U.S. Department of Energy Joint Genome Institute, a DOE Office of Science User Facility, is supported under Contract No. DE-AC02-05CH11231.

## Author Contributions

Conceptualization, J.D., F.B.Y., and S.Q.; Methodology, J.D., F.B.Y., and S.Q.; Software, J.D. and F.B.Y.; Investigation, J.D., F.B.Y., and S.Q.; Writing, J.D., F.B.Y., and S.Q.; Supervision, F.B.Y. and S.Q.

## Data Availability

Annotated contigs are available through JGI’s IMG portal https://img.jgi.doe.gov/mer/under IMG Genome IDs 3300006068 and 3300006065.

Sequencing data is available under NCBI SRA under BioProject PRJNA378813. PhaMers software is be available at https://github.com/jondeaton/phamers

